# Male parental investment reflects the level of partner contributions and brood value in tree swallows

**DOI:** 10.1101/216119

**Authors:** Ádám Z. Lendvai, Çağlar Akçay, Mark Stanback, Mark F. Haussmann, Ignacio T. Moore, Frances Bonier

## Abstract

Biparental care presents an interesting case of cooperation and conflict between unrelated individuals. Several models have been proposed to explain how parents should respond to changes in each other’s parental care to maximize their own fitness, predicting no change, partial compensation, or matching effort as a response. Here, we present an experiment in tree swallows (*Tachycineta bicolor*) in which we increased the parental care of females by presenting them, but not their mates, with additional nestling begging calls using automated playbacks. We performed this experiment in two populations differing in future breeding opportunities and thus the intensity of conflict over current parental care. We found that in response to a temporary increase in female parental effort, males in the northern population with lower sexual conflict matched the increased effort, whereas males in the southern population did not. We also found that increases in parental care during playbacks were driven by the females (i.e., females initiated the increased effort and their mates followed them) in the northern population but not the southern population. These results support the idea that with incomplete information about the brood value and need, cues or signals from the partner might become important in coordinating parental care.

## Introduction

In many animal species, parents provide some form of parental care before offspring become independent [1]. Although in many species parental care is only provided by a single parent, biparental care is especially common among birds [2]. Biparental care provides an interesting case study in cooperation between unrelated individuals as well as conflict between sexes [3–5], given parental care is often costly and reduces the chances of future breeding [6]. Although both parents gain a fitness benefit from increased parental care via increased survival of the offspring, each parent is better off if the other parent supplies the majority of the care [6–8]. This sexual conflict may be particularly severe in cases where parents have good future reproduction opportunities and investing less into current reproduction benefits their chances of future reproduction. On the flip side, sexual conflict may be low or absent if future reproductive opportunities for the parents are limited as lower investment would not necessarily translate into improved reproduction later.

Several types of models have been proposed to predict how conflict over parental care may be resolved. These models differ in the assumptions they make regarding behavioural strategies available to parents and consequently the predictions they offer when a parent increases or decreases parental care. The first type of model is the ‘sealed-bid model’ [7] which assumes that parents will be behaviourally insensitive to changes in each other’s reproductive effort, instead engaging in a fixed level of parental effort [7,9]. In this model, changes in parental effort of one of the sexes only occurs through evolutionary change. An alternative prediction comes from ‘negotiation models’ that assume that parents are able to respond to changes in partner investment. In these models, parents are viewed as being in partial conflict over the amount of parental care they provide, with each parent preferring the other parent do more, and themselves do less. When one parent decreases their effort, the other parent is predicted to compensate by increasing their own effort but this compensation has to be incomplete for this model to be evolutionarily stable [10–14]. The logical extension of these models is that when a parent increases parental effort, the other parent should compensate by decreasing their own parental effort [15]. A third model, the ‘information model’ [4,16] assumes that parents have only incomplete information about the brood value and need, and they use their mate’s parental effort as a cue in determining the level of parental care needed to optimize reproductive success. Under this model, parents are expected to match an increase in the other parent’s effort with an increase in their own effort. Finally, the ‘perfect family’ model [17] also predicts a matching response when one parent increases their parental effort, but in this model (unlike any other above), the parents are assumed to be able to communicate directly to coordinate their parental efforts.

Several studies have tested how the change in one parents’ provisioning behaviour affects the parental behaviour of its partner. In a meta-analysis, the general pattern across these studies was partial compensation, consistent with the predictions of the negotiation models and the information model when information about brood value is complete and symmetrical [5]. However, most of the experimental tests used handicapping, i.e. an experimentally-induced decrease in one parent’s parental effort (e.g. by feather clipping, weight addition, or hormonal manipulation), or mate removal [5]. Both handicapping and mate removal may have limitations in addressing models to explain parental care. Handicapping not only alters the focal parent’s contribution to offspring care, but also potentially its physical appearance and/or attractiveness, which may confound how its partner will interpret the treatment [18]. Mate removal simulates the desertion or death of one parent, and the response of the remaining parent may be contingent on the complete absence of the partner, instead of representative of responses to changes in ongoing biparental care [5,12,14,19]. Only a few studies have sought to experimentally increase the parental effort of one parent. Two of these studies [13,20] found evidence for matching of partner effort while one [15] found partial compensation (i.e. decrease of own effort in response to increase in partner effort).

In this study, we used playbacks of recordings of nestling begging calls as an experimental treatment to stimulate an increase in parental care (nestling provisioning) in female tree swallows (*Tachycineta bicolor)* and then measured the response of their mates. Previous studies that used begging calls used only short-term modification of parental behaviour (usually a 1h-playback session) [16,20]. We designed an automated broadcasting and recording system [21] that allowed us to deliver the experimental stimulus only to focal females and the treatment lasted up to six hours during the day of the study. We have previously shown that this experimental manipulation temporarily increased the females’ feeding rate [22]. Here, we investigated how their partners reacted to the manipulation and tested predictions from the hypotheses presented above. The sealed-bid model predicts that males will not be responsive to short-term changes in female provisioning behaviour. The negotiation models predict that when females increase their provisioning rate, the males should decrease their own effort. Finally, the information and perfect family models predict that males should match the female response to the playback, thus increasing their own parental effort.

We also tested the hypothesis that the resolution of the sexual conflict may depend on the value of the current reproduction. Brood value is a concept that combines several life-history traits to summarize the value of the current brood relative to the potential for future reproduction [23,24]. In populations with lower potential for future reproduction (due to a shorter breeding season, higher adult mortality rates, etc.) the current brood is more valuable compared to a brood of the same size in a population where the probability of future reproduction is higher. Brood value therefore is related to the intensity of sexual conflict and is likely to affect negotiation rules that parents use in determining their response to changes in partner behaviour. We examined the role of brood value by carrying out the experiment in two populations of tree swallows that differ in brood value. One population, in Ontario, Canada, has lower annual survival rates, a shorter breeding season, and thus higher brood value and lower sexual conflict. The other population, in North Carolina, USA, has higher annual survival rates and a longer breeding season that allows some birds to successfully nest twice in one year, and thus lowers the current brood value and increases sexual conflict [22]. Because of these differences in brood value, we predicted that male tree swallows in the Ontario population would be more likely to match their partner’s increased parental effort than the males in North Carolina.

## Methods

### Study site and species

We studied tree swallows at two field sites where swallows nest in artificial nest boxes: Queen’s University Biological Station, Ontario, Canada (N44°34’2”, W76°19’26”, 121 m elevation) and Davidson College, Davidson, North Carolina, USA (N34°31’32”, W80°52’40”, 240 m elevation). These two sites differ in the length of the breeding season (longer seasons in North Carolina) and annual survival rates (higher in North Carolina). Tree swallows have high breeding site fidelity, and so return to the breeding population is often used as a proxy of annual survival [25]. In the North Carolina population, return rates are around 50% for females (51% in 2015), which is higher than in the Ontario population (average 22% between 1975-2012, range 10-45%). Similar findings have been reported in other studies comparing southern and northern populations of tree swallows [26].

### Subjects

We captured birds using box traps at their nest or placing our hands over the nest entrance. We caught females on day 10 of the incubation period and males on day 2 or 3 post-hatching. We recorded body measurements (tarsus, wing chord, weight, skull size) and marked birds with a numbered metal leg band (US Fish and Wildlife Service or Canadian Wildlife Service) and a unique passive integrated transponder (PIT) tag that was integrated into a coloured (red for females, blue for males) plastic leg band (EM4102 tags from IB Technology, UK). More details on the field methods can be found in [22].

### Playback experiment

Nestling begging calls were recorded as described in [22]. Briefly, we recorded the calls from 10 nests on day 6 post-hatching by tapping at the nest entrance to simulate the sound of an arriving parent and pointing a directional microphone (Sennheiser ME66/K6 directional microphone connected to a Marantz PMD 660 solid-state recorder) into the nest. We used the software Syrinx (John Burt, Seattle, WA; www.syrinxpc.com) to create 30 second stimulus files from the recordings thus obtained, as described in [22]. The initial calling rate was ~14 begs/sec (consisting of overlapping calls by multiple nestlings) that gradually decreased to a constant ~4 begs/sec (see supplementary material for an example stimulus). The 10 stimulus files were randomly allocated to the treatment nests.

### Playback set-up

We used a radio-frequency identification (RFID) reader attached to a micro-computer (Raspberry PI) to carry out the playbacks automatically [21]. The computer was programmed to carry out playbacks every time the female (but not the male) was perched at the nest entrance (where the RFID antenna was attached). Each playback lasted 30 seconds after the RFID reader detected the female’s PIT tag, and there was a refractory period of 2 minutes from the start of each playback (to avoid situations where the playback would be triggered by the female leaving the nest soon after she had entered). We used earbud headphones (Sony MDRE9LP, Sony Inc.) secured with tape at the back of the nest box as speakers [see for picture:, 21], playing calls at approximately 55 dB (measured from approximately 10 cm), which is comparable to call amplitudes of tree swallow nestlings at that age [27]. The playback apparatus was also installed in control nests, but no sound was played. Treatments were allocated to the nests using a randomized block design, to control for seasonal differences. In order to assess the possible effects of the playback on the nestlings’ begging calls, we also recorded nestling begging calls in 9 nests in Ontario following the methodology described in [28].

We set-up the playback systems at around 7am on day 6 post hatching and playbacks stopped approximately 6 hours later, after which we captured the females to obtain a blood sample for hormone analysis [22]. We had 20 control and 16 playback nests in NC and 12 control and 12 playback nests in Ontario.

### Quantifying parental effort

We quantified rates of parental visits to the nest, used here as the measure of parental effort, in two ways: first, we carried out 1-hour direct observations on day 5 and day 6 (the day before and the day of the treatments). For these, an observer sat ~30 m from the nest and noted every visit of the male and female using a spotting scope and a voice recorder. We also quantified visit rates from the RFID records as described in detail in [29]. We checked the visit rates from 1-hour nest watches against the visit rates calculated from RFID logs of the same time periods. There was a high correspondence between the two measures [29]. Because the RFID observations spanned the entire duration of the experiment we used these data as the primary measure of parental visit rates. Visit rates are an excellent measure of the feeding rates in the tree swallows, as most visits are for feeding [30].

### Statistical analyses

We used generalized linear mixed models (GLMM) to assess the effects of treatment and population on male behavioural data. For all models, we first fitted a fully parameterized model with all interactions included and then used a model selection based on the Akaike Information Criterion (AICc). We performed model averaging of the best models (within 2 AICc units compared with the model with the lowest AICc value) and report the model averaged coefficients. The conclusions drawn from this model selection procedure were consistent with an alternative approach using stepwise model selection and achieving a single, minimal adequate model. We analysed the feeding rates derived from RFID recordings with GLMMs using the fixed factors treatment (playback vs. control), population (Ontario vs. North Carolina), time period, and their interactions. For the time period factor, initially we used four levels: pre-treatment (day 5) feeding rates (6 hrs. during the same time of day as the experimental period on the next day) and feeding rates from the period while the playback or control treatment was in effect in day 6, which we further divided into three two-hour periods to assess any temporal changes in effects of playback on female behaviour. We included playback stimulus recording and bird ID as random factors and also included an offset variable for log of duration of playback to control for variation in how long the birds were exposed to the playbacks (mean= 6.24 ± 0.05 SE hours).

In a next step, we analysed the *change* in male visit rate from the pre-treatment period (day 5) to the experimental period (day 6) as a function of the treatment, population, and the change in provisioning behaviour of females over the same time period. The change in visit rate was calculated as the difference in the number of feeding visits per time between day 5 and day 6, where time was the total 6 hours of experimental period on day 6, and the same time frame on day 5. Because in this case, we only had one male and one female change in visit rate value per nests, this was analysed by a GLM with male change in visit rate as the response variable and with female change in visit rate, treatment, and population as explanatory variables. The initial model contained all possible interactions.

Finally, we analysed the relationship between male and female feeding rate over a more continuous scale using time series analyses. To do that, the event recordings from the RFID logs were transformed into time series using moving averages (calculating over a frame of 60 minutes and in 20 minutes steps). We tested various combinations of these parameters, and they gave qualitatively similar results (i.e., did not change our conclusions). During the time series analyses, first we tested if the male and the female time series are significantly related to each other using cross correlation analyses within ± 80 minutes (that corresponds to lag = 4). In those pairs where we found a significant relationship between the time series, we determined the peak of the cross-correlation distribution, and the lag that corresponded to it. Finally, we repeated these analyses using only the experimental period to test how the experimentally induced change in female behaviour may affect their partners’ parental contributions.

## Results

### Feeding rates

The playback had a transient (first 2 hours) and positive effect on the feeding rate of females in both populations (Table 1).

**Table 1:**
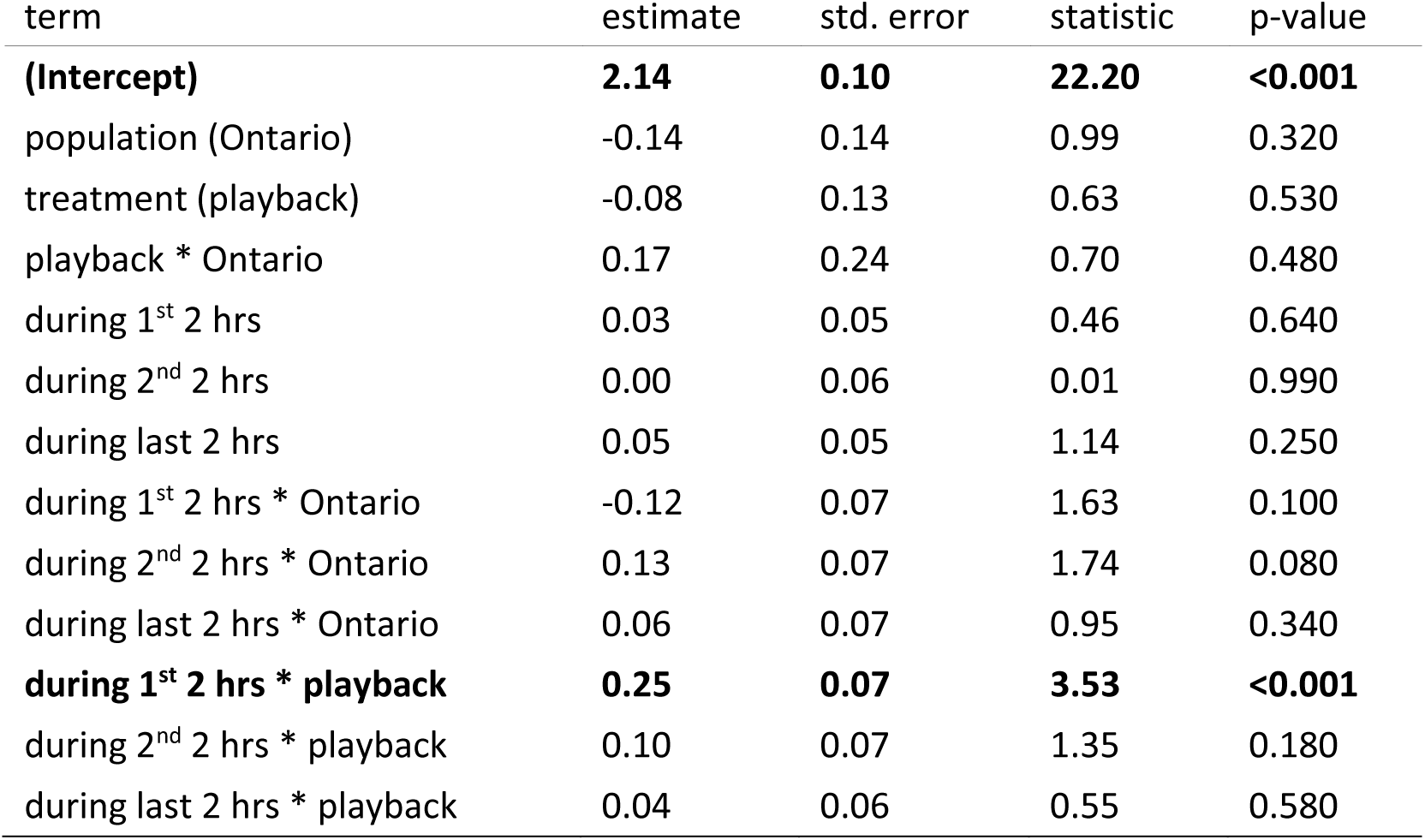
Model averaged parameter estimates of a General Linear Mixed-effects Model (Poisson error and log-link) on the effects of playback on female feeding rate. The intercept corresponds to the baseline level (pre-treatment, day5), North Carolina, and control treatment.

| term | estimate | std. error | statistic | p-value |
| --- | --- | --- | --- | --- |
| **(Intercept)** | **2.14** | **0.10** | **22.20** | **<0.001** |
| population (Ontario) | -0.14 | 0.14 | 0.99 | 0.320 |
| treatment (playback) | -0.08 | 0.13 | 0.63 | 0.530 |
| playback * Ontario | 0.17 | 0.24 | 0.70 | 0.480 |
| during 1<sup>st</sup> 2 hrs | 0.03 | 0.05 | 0.46 | 0.640 |
| during 2<sup>nd</sup> 2 hrs | 0.00 | 0.06 | 0.01 | 0.990 |
| during last 2 hrs | 0.05 | 0.05 | 1.14 | 0.250 |
| during 1<sup>st</sup> 2 hrs * Ontario | -0.12 | 0.07 | 1.63 | 0.100 |
| during 2<sup>nd</sup> 2 hrs * Ontario | 0.13 | 0.07 | 1.74 | 0.080 |
| during last 2 hrs * Ontario | 0.06 | 0.07 | 0.95 | 0.340 |
| **during 1<sup>st</sup> 2 hrs \* playback** | **0.25** | **0.07** | **3.53** | **<0.001** |
| during 2<sup>nd</sup> 2 hrs * playback | 0.10 | 0.07 | 1.35 | 0.180 |
| during last 2 hrs * playback | 0.04 | 0.06 | 0.55 | 0.580 |

We tested how males reacted to the change in behaviour of their mates during the same periods, and we found a significant 3-way interaction (treatment*population*time period), indicating that the males’ behaviour was affected by the playbacks broadcast to their partner, and this response was different between the two populations (Table 2). Specifically, this model showed that in control nests, during the first two hours of treatment, males tended to decrease their feeding rate compared to day 5, but this effect was more pronounced in Ontario. The decrease in parental care in the first two hours of controls is probably due to the disturbance caused by setting up the dummy playback apparatus, which disappeared by the second hour of playback and resulted in higher feeding rates in Ontario than in North Carolina However, in playback nests, this initial decrease is not seen in Ontario, on the contrary, during the first two hours of the playback, males increased their feeding rate, and increased it further in the second two-hour period (Fig. 1.)

**Table 2:**
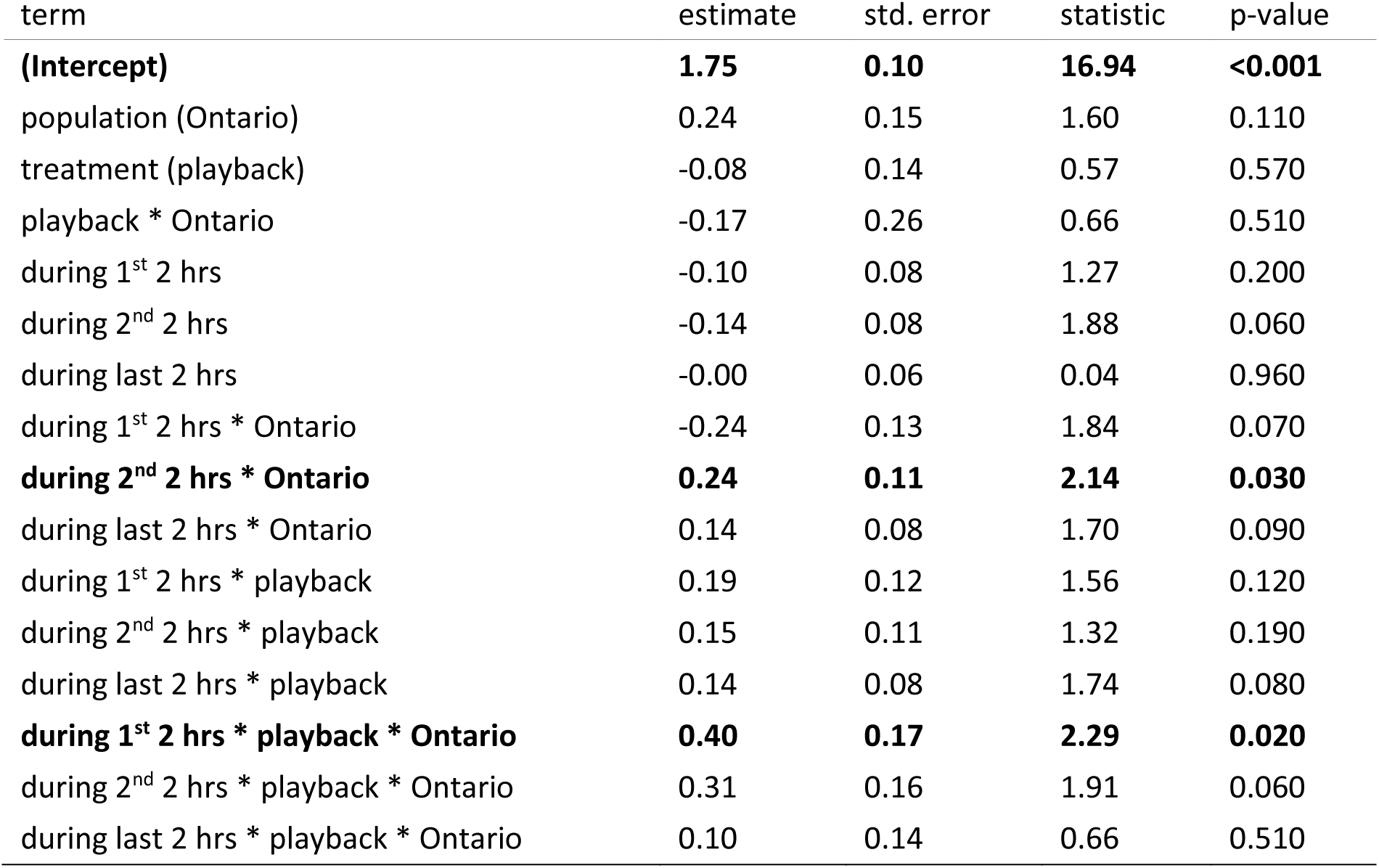
Model averaged parameter estimates of a General Linear Mixed-effects Model (Poisson error and log-link) on the effects of playback on male feeding rate. The intercept corresponds to the baseline level (pre-treatment, day5), North Carolina, and control treatment.

**Figure 1:**
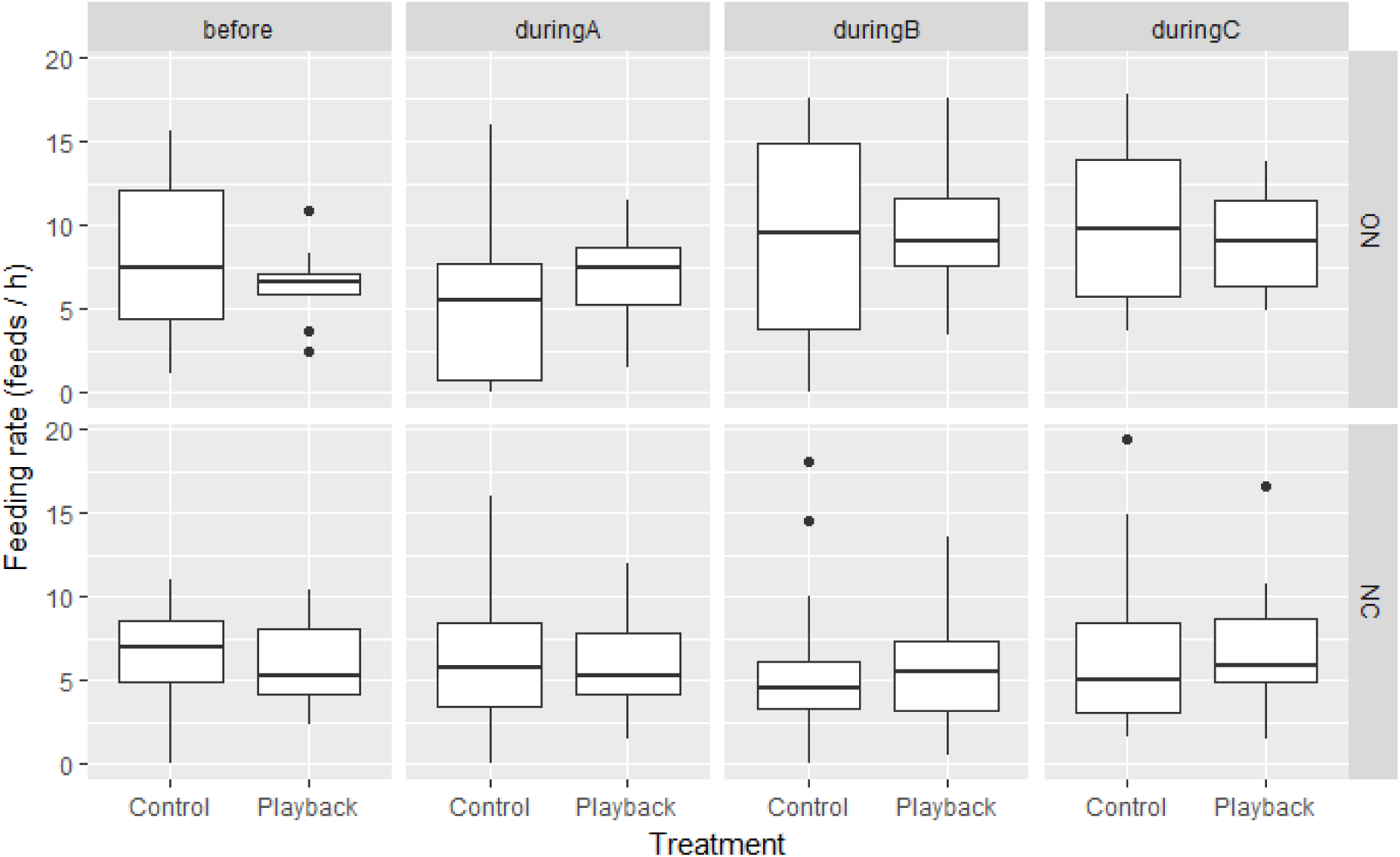
Nestling provisioning behaviour in male tree swallows in two populations (NC = North Carolina, USA; ON = Ontario, Canada) before and during broadcasting experimental nestling begging stimuli for female parents. The experimental period was divided into three 2 hours subsets (A, B, C).

### Change in feeding rates

Analysing the change in feeding rate from the baseline (day 5) to the entire experimental period (day 6) revealed that the most important predictor of how males changed their behaviour was the change of their partners’ provisioning rate (Fig. 2, Table 3). In control nests, most pairs’ behaviour followed the predictions of the matching hypothesis, i.e. an increase or a reduction in female feeding rate was mirrored by a similar change in the male’s behaviour. This pattern remained the most common one in response to the playback as well. Interestingly, males in Ontario showed a stronger response to the playback treatment than their mates (even though playback was only broadcast to females): despite the modest increase in female feeding rate during the playback period, all but one male in Ontario increased their feeding rate compared to the baseline (Fig. 2b) resulting in a significant increase in that population (Fig. 2d).

**Figure 2:**
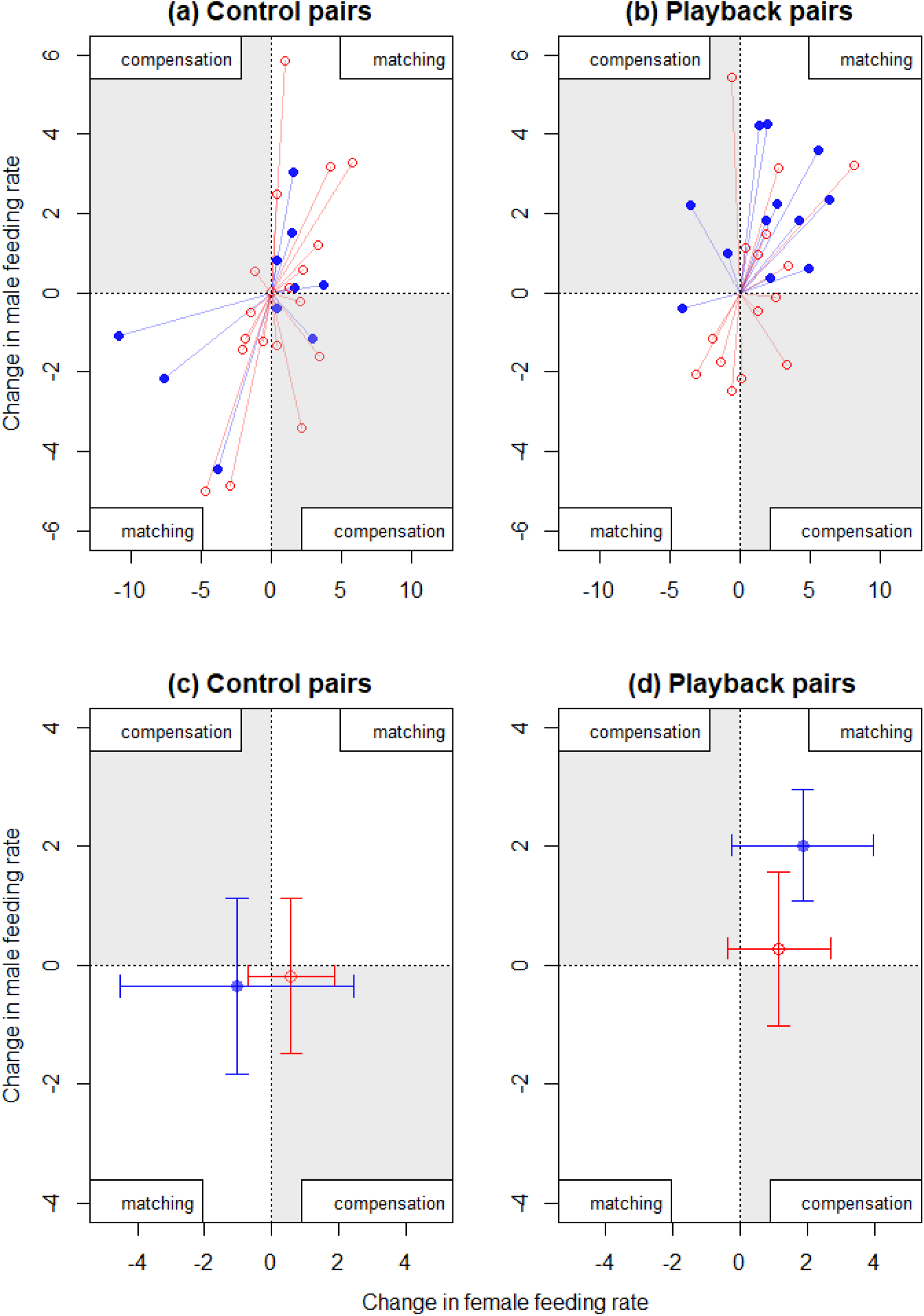
Relationship between changes in feeding rates from pre-treatment (day 5) to the experimental periods (day 6) within tree swallow pairs. Panel (a) and (b) shows the individual data points, panel (c) and (d) shows the means and 95% confidence intervals. Red colour and open circles indicates North Carolina, blue colour and filled circles denotes Ontario.

**Table 3:** Model averaged parameter estimates for the model analysing change in feeding rate from the baseline (day 5) to the experimental period (day 6).

| term | estimate | std. error | statistic | p-value |
| --- | --- | --- | --- | --- |
| (Intercept) | -0.46 | 0.43 | 1.05 | 0.294 |
| **change in female feeding rate** | **0.41** | **0.13** | **3.10** | **0.002** |
| treatment (playback) | 0.74 | 0.56 | 1.29 | 0.197 |
| population (Ontario) | 1.06 | 0.57 | 1.80 | 0.072 |
| change in female feeding rate * Ontario | -0.24 | 0.17 | 1.37 | 0.169 |

### Pair coordination

In many pairs the two parents seem to follow each other’s behaviour closely (Fig. S2). In both populations, most pairs show a matching pattern (i.e. the cross-correlation coefficient before the treatment was significantly positive in 27 out of 36 nests in North Carolina and 19 out of 24 nests in Ontario, Figure S3). This pattern was unaffected by the treatment or the experimental period (day 5 vs. day 6). As a result, the analyses of the cross-correlation coefficients showed that the best model was the null-model (ΔAIC from the next model was > 4), where only the intercept was significantly positive confirming the positive relationship between the parents’ provisioning efforts (Figure 3, Table 4). Thus, increases or decreases in one parent’s provisioning effort tended to be matched by the other parent’s provisioning effort in the same direction irrespective of the treatment and experimental period.

**Figure 3:**
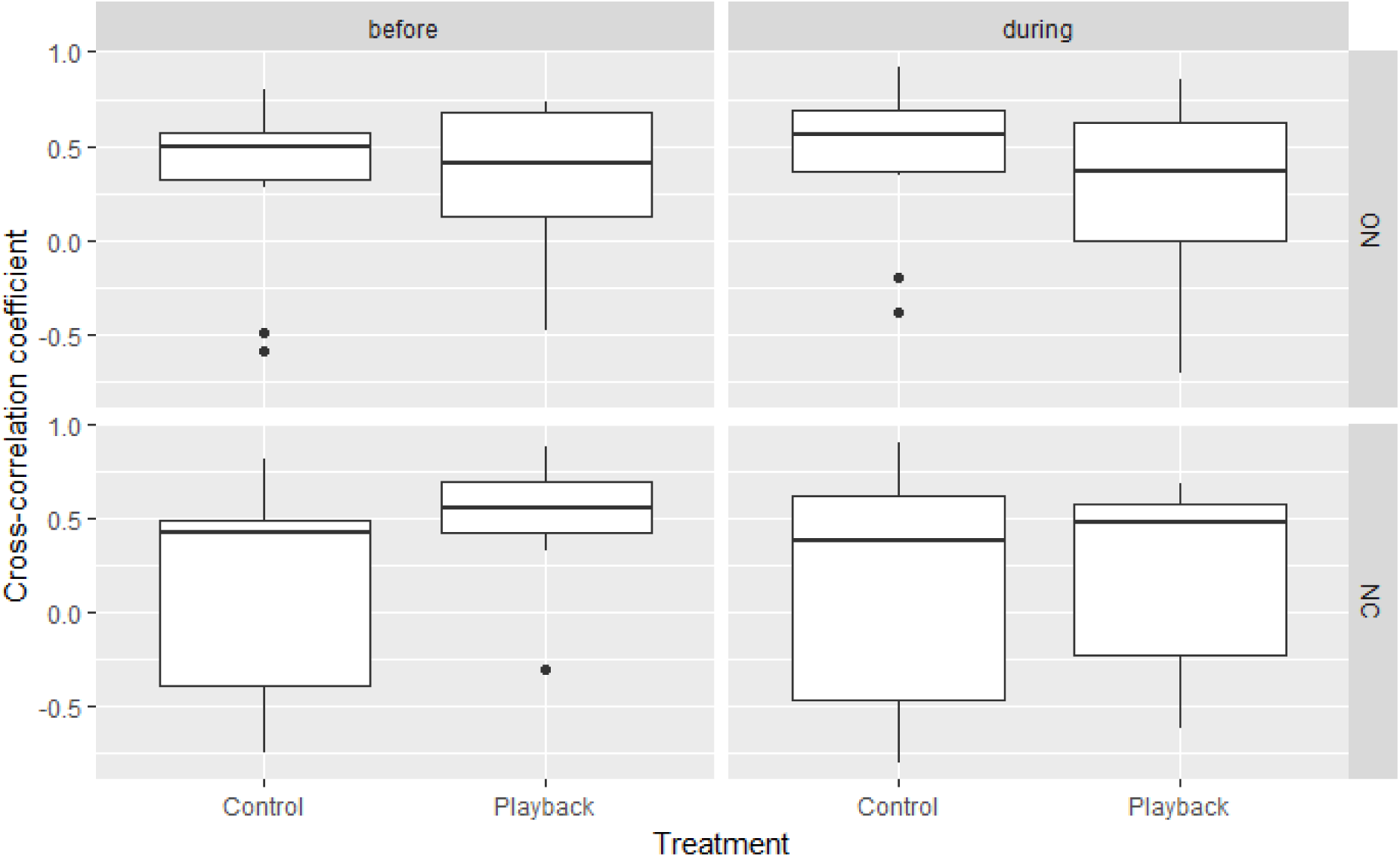
Parental coordination (cross-correlation coefficients) for tree swallow pairs in two populations (Ontario: ON and North Carolina: NC). Positive values indicate a matching response (an increase in provisioning in one sex results in an increase of its partner), while negative values indicate a compensation response.

**Table 4:** Model averaged parameter estimates for the model analysing peak cross-correlation coefficients between male and female feeding rate.

| term | estimate | std. error | statistic | p-value |
| --- | --- | --- | --- | --- |
| **(Intercept)** | **0.28** | **0.05** | **5.10** | **<0.001** |
| treatment (playback) | 0.10 | 0.09 | 1.07 | 0.283 |
| population (Ontario) | 0.10 | 0.09 | 1.04 | 0.297 |
| period (day 6 – during playback) | -0.08 | 0.09 | 0.87 | 0.382 |

However, treatment had a modest effect on who was leading the changes in provisioning behaviour (as measured by the lag at which the cross-correlation coefficients were maximal with positive lags indicating female-led changes and negative lags indicating male-led changes, Figure S3), but only in the Ontario population, resulting in a marginally non-significant three-way interaction (Table 5). While changes in provisioning behaviour was highly synchronous among pair members in NC during the pre-manipulation period (day 5) (the lag corresponding to the peak cross-correlation was close to zero), in Ontario, females tended to follow changes in their partners’ behaviour before the experimental treatment. This pattern changed in the Ontario population in response to the treatment: in the playback group variation in the feeding rates became female-driven in most nests with males following the change in female provisioning, i.e. the lags became positive (Figure 4, Table 5). The control group, by contrast, became more synchronous (with the lag becoming near zero).

**Table 5:** Model averaged parameter estimates for the model analysing the temporal association between male and female feeding rates.

| term | estimate | std. error | statistic | p-value |
| --- | --- | --- | --- | --- |
| (Intercept) | 0.02 | 0.37 | 0.06 | 0.955 |
| population (Ontario) | -0.53 | 0.76 | 0.68 | 0.494 |
| period (day 6- during playback) | 0.14 | 0.69 | 0.20 | 0.839 |
| treatment (Playback) | 0.39 | 0.74 | 0.52 | 0.601 |
| Ontario * day 6 | 0.32 | 1.36 | 0.24 | 0.814 |
| Ontario * Playback | -1.15 | 1.26 | 0.90 | 0.367 |
| Playback * day 6 | -1.47 | 1.13 | 1.29 | 0.195 |
| Ontario * Playback * day 6 | 3.39 | 1.78 | 1.89 | 0.059 |

**Figure 4:**
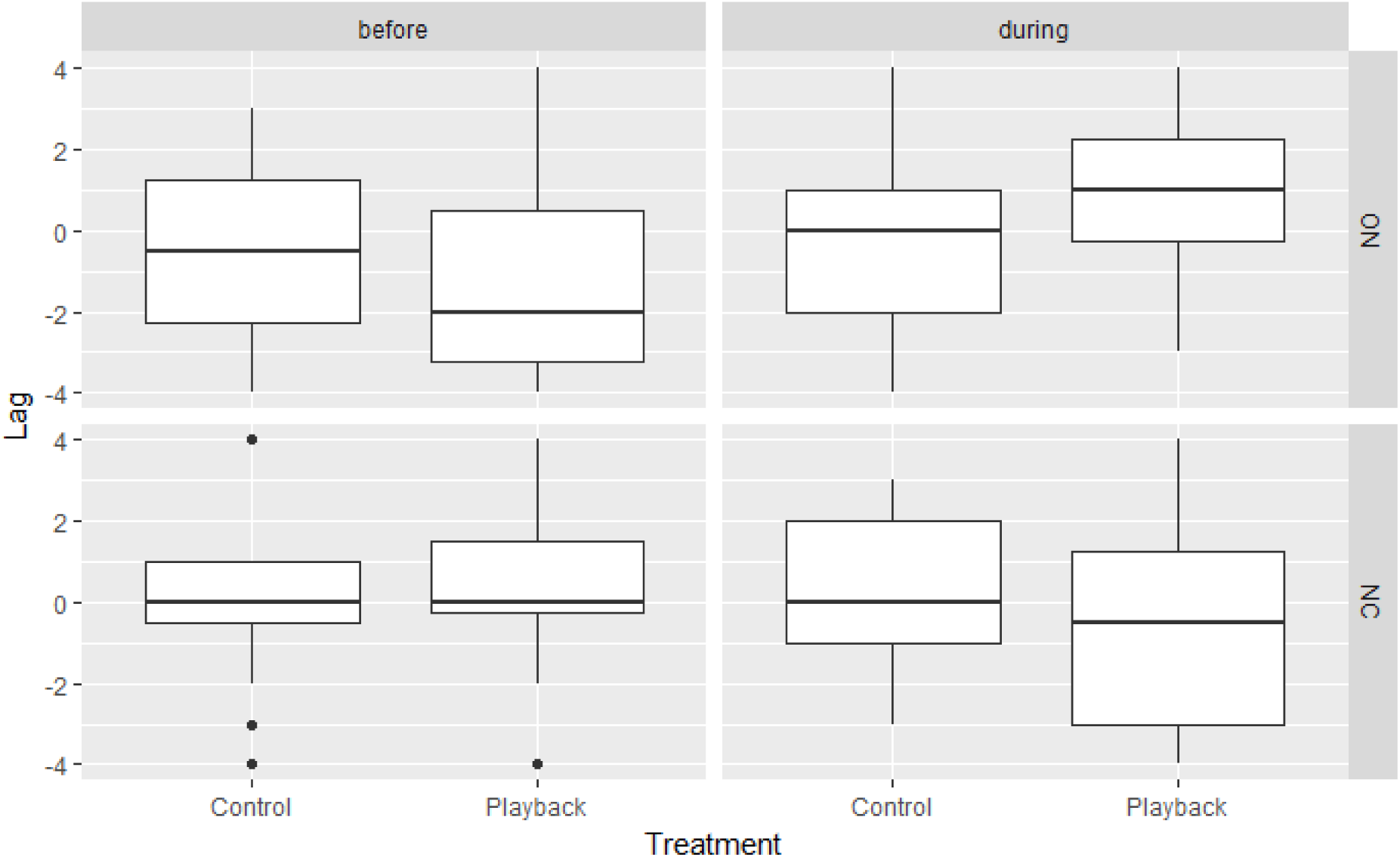
Temporal association between provisioning behaviour of tree swallow parents in two populations (Ontario: ON and North Carolina: NC) as shown by lags at which the cross-correlation is maximal between female and male feeding rate. The closer the lag is to zero, the more synchronous the parents are. Positive lags indicate that the male’s parental behaviour follows the female’s behaviour; negative lags mean that female behaviour follows the male’s behaviour.

## Discussion

We found that when experimental nestling begging calls were broadcast to female tree swallows, males in two different populations, differing in latitude and brood value, showed marked differences in their reaction. Males in Ontario significantly increased their feeding rates compared to the previous day, whereas control males did not. In contrast, males in North Carolina did not show any increase in their feeding rates, despite the fact that females in both populations showed a transient increase in their feeding rates [22]. The finding that males in the Ontario population responded with an increase in feeding rates is consistent with the predictions of the information model [4] and the perfect family model of biparental care [17]. The most likely explanation for this result is that males adjusted their parental effort based on cues or signals from their mates.

The effect of begging playbacks on male feeding rates suggests that males matched increases in female feeding rates, at least in the Ontario population. Interestingly, although the direction of the males’ response matched that of their mates, the magnitude of their response was actually larger. Time series analysis also showed that, at least in some pairs, males substantially increased their feeding rate during the playback period before the increase of their mates’ contribution was apparent.

The mechanism by which male parents match females is currently unknown, but there are several plausible explanations, which are not mutually exclusive. First, males may have been simply responding directly to offspring begging call playbacks they overheard. This could occur if they were perched at the nest box entrance or on the box at the time of a playback. We designed the playback system to minimize this possibility: after the initial playback that was played upon the arrival of the female, there was a refractory period of two minutes, so during most feeding visits, when females left the nest after entering to feed, a playback was not played. We also verified with the RFID data that it was very rare for males to be perched at the box entrance while the playback was active and the female was inside the box (see Supplementary Information). Additionally, behavioural observations suggest that males were rarely present on the box when females triggered playbacks and the latter measure did not differ between treatment groups or experimental periods (day 5 vs. day 6 – see Supplementary Information). Thus, it is unlikely that males directly overheard the playbacks.

A second possibility is that the playbacks may alter nestling begging behaviour during male visits. In particular, nestlings may anticipate higher acoustic competition from their real and simulated nest mates [31], and therefore increase their own begging rate, which has been shown to influence parental feeding in this species [32]. There is evidence that nestlings tend to beg more intensely in larger broods [33], which suggests that acoustic competition potentially could lead to higher calling rates, even when there is no playback (i.e. during the male’s visit). We recorded nestling begging behaviour in a subset of nests in Ontario, and the analysis of these audio recordings showed that while begging rates increased from day 5 to day 6, this increase was similar in control and playback nests (see Supplementary Information). An earlier study with great tits testing this hypothesis also found that begging behaviours were not effective in predicting partner response [13].

A third possibility is that males may match their feeding rate to that of the females using a tit-for-tat style alternation of provisioning where individuals time their feeding according to their partners’ provisioning [34,35]. Under this scenario we would expect that because the treatment is targeted to females, they are the ones who initially start increasing their feeding rate and the males match their effort. This explanation is consistent with the results of the time series analysis, where we observed a change in the temporal association in the experimental group in Ontario. While the cross-correlation of the male and female feeding rates was significantly positive, and did not change in response to the treatment, the lag in the playback group in the Ontario population shifted from slightly male-driven towards slightly female-driven. Note that if the parents are well coordinated, we do not expect a large change in time lags, because that would suggest that there is a considerable delay in how parents respond to the changes in their partner’s behaviour. A moderate shift towards a more female-driven cross-correlation is consistent with the explanation that males were responding to cues provided by the females.

Males may have adjusted their feeding effort as a result of communication with the female receiving the playbacks [17,36,37]. This explanation would be most consistent with the perfect family hypothesis wherein males adjust their parental effort based on cues or signals from their mates. It could also explain the result that in some pairs the increase in male feeding rate precedes that of the females: if the females can communicate the need of a higher contribution towards their partner early during the experimental period, then some males may adjust their provisioning behaviour sooner than their partner.

While a matching response is consistent with both the perfect family and the information model, it is also important to note that the predictions of the information model depend on the cues available to the parents. This model, contrary to the perfect family model, does not assume direct communication between the parents and only predicts a matching response if the parents do not have symmetrical and complete information about the needs of the offspring, which might be the case in chick-rearing. In this experiment we deliberately created conflicting information available to the males. While cues coming from the nestlings did not convey the message of hungry chicks (nestling begging rate did not seem to be affected by the treatment -see Supplementary Material), the females may have communicated a need for higher level of provisioning. If the information model is correct, our results suggest that the males use cues from their partners (either direct information through communication as the perfect family hypothesis suggests, or indirect cues, such as their feeding rate). This may also explain why another study, that also set out to increase (rather than to decrease) one parent’s effort found the opposite result, i.e. that the predominant response of focal parent’s mates was reversible compensation and not matching [15]. The latter study was carried out during incubation, and both parents could easily detect the difference between optimal and actual egg temperature and use this as a direct cue in the decision about parental care [15].

In contrast to the findings in Ontario, there was no effect of playback on male behaviour in the North Carolina population. This finding is consistent with the brood value hypothesis as the current brood in the Ontario population should be more valuable to the males compared to the broods in North Carolina, where adult survival is higher and the reproductive season is longer. Interestingly however, we did not see a population difference in females in our earlier analyses [22], so the brood value hypothesis only seems to hold for males but not females. This difference between the sexes could be due to the fact that brood values between the populations may vary less for females than for males. The brood value for males may vary depending on the extra-pair paternity [38] or sex-specific differences in adult annual survival rates. It is currently not known how extra-pair paternity varies with latitude and life history in this species.

Taken together, our results are most consistent with the perfect family and the information model of biparental care and suggest that males may use cues from their partners to decide about their actual parental effort. The intriguing possibility that parents can directly convey such messages to their partners through communication supports the notion that animal communication signals may be much more complex than previously thought [36,39].

## Acknowledgments

We are grateful to Alice Domalik, and Pria St John (Queen’s University), and Drew Gill and Spencer Gill (Davidson College) for excellent help in the field and for Fruzsina Demcsák (University of Debrecen) for analysing the begging recordings.

## Data accessibility

All data are available as part of the Supplementary Material.

## Animal Ethics

We confirm that the procedures used in the study followed the guidelines for animal care outlined by Animal Behaviour Society and Association for the Study of Animal Behaviour, and were approved by approved by the Institutional Animal Care and Use Committee at Virginia Tech (#12–020) and Animal Care Committee of Queen’s University (#2013-019) and the Canadian Wildlife Service (#10771).

## Permission to carry out fieldwork

The field research was conducted with a permit from US Geological Survey Bird Banding Laboratory to MS (#22742) and Canadian Wildlife Service permit to FB (#10771)

## Author contributions

ÇA and ÁZL designed and coordinated the study, collected field data, carried out data analysis and drafted the manuscript, MS collected field data and helped draft the manuscript, MFH contributed to the design of the study and helped draft the manuscript, FB collected field data, contributed to the design of the study and helped draft the manuscript, ITM contributed to the design of the study and helped draft the manuscript. All authors gave final approval for publication.

## Supplementary material

Original data is accessible as a Supplementary file.

**Figure S1:**
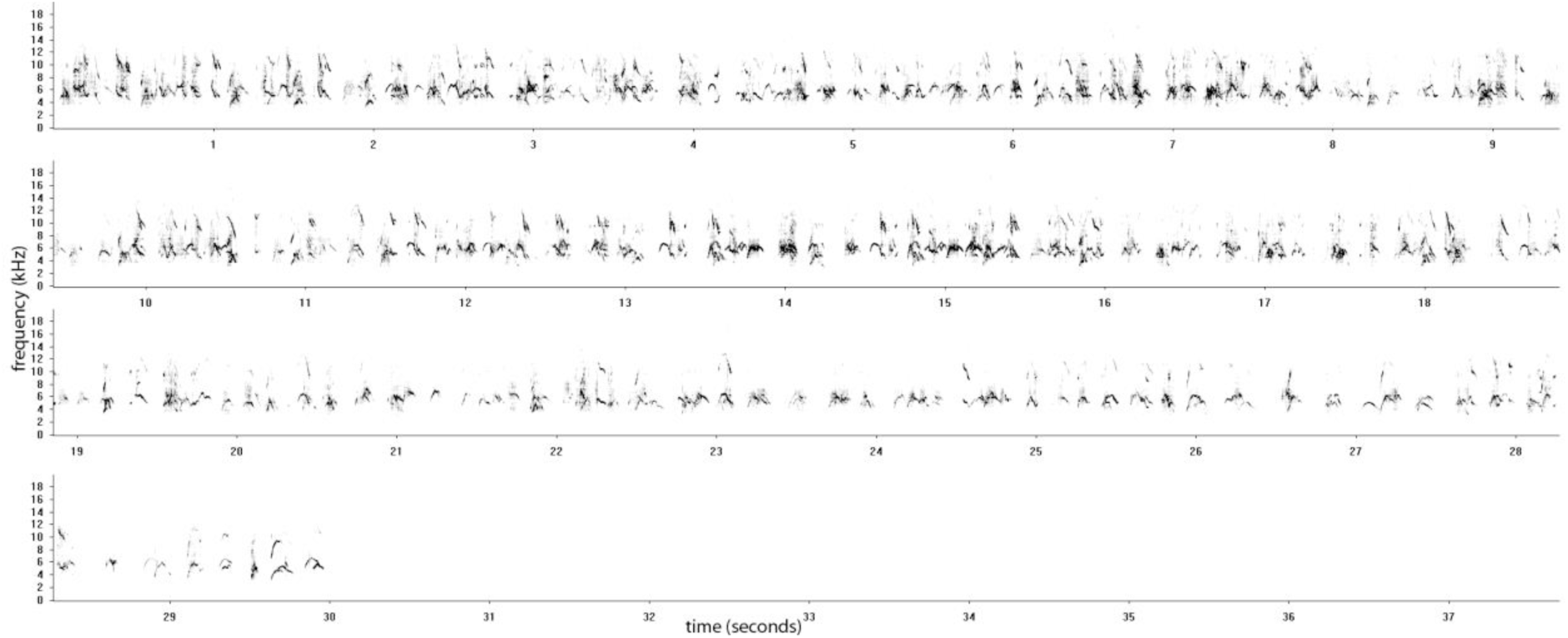
An example sonogram of a stimulus file used in the experiment.

**Figure S2:**
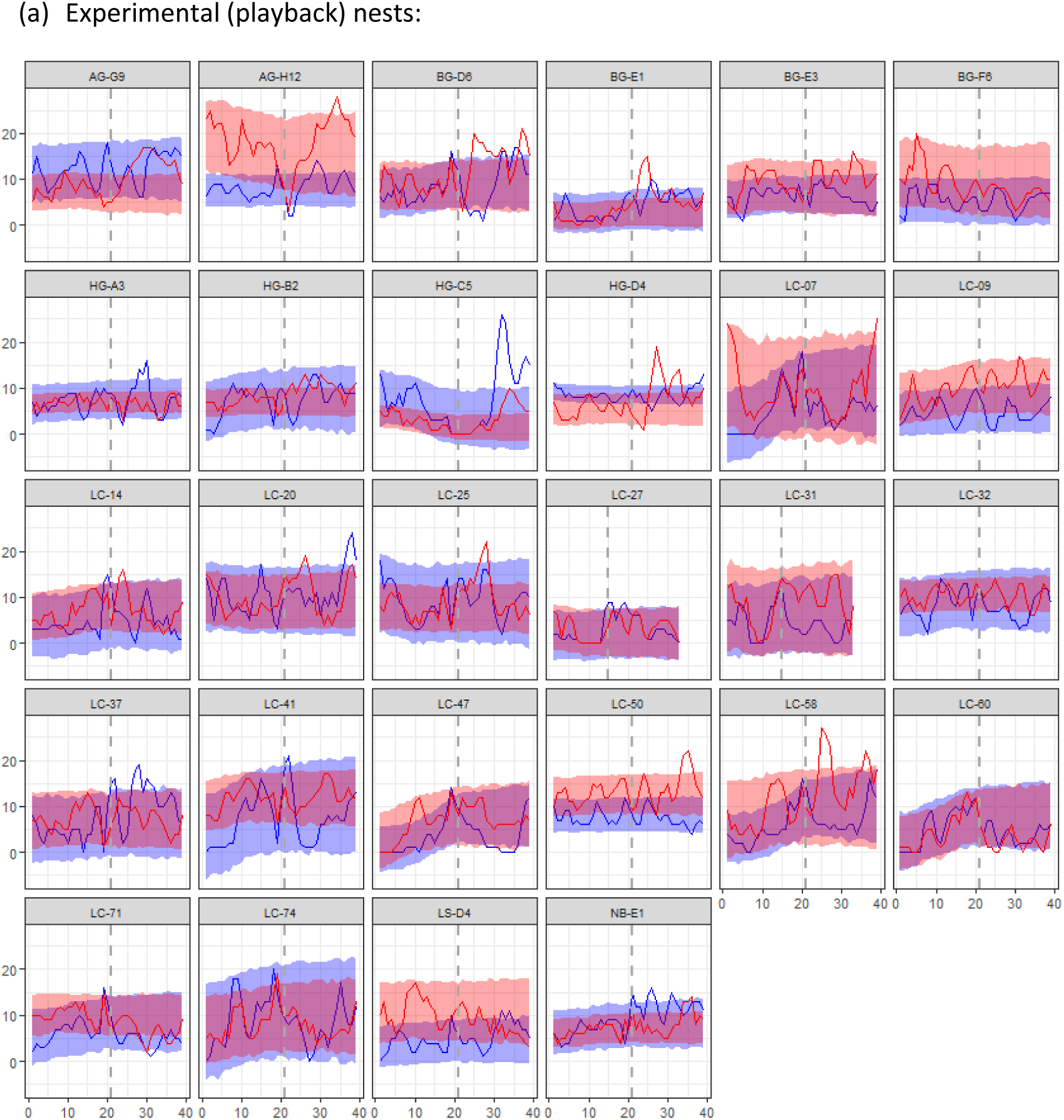
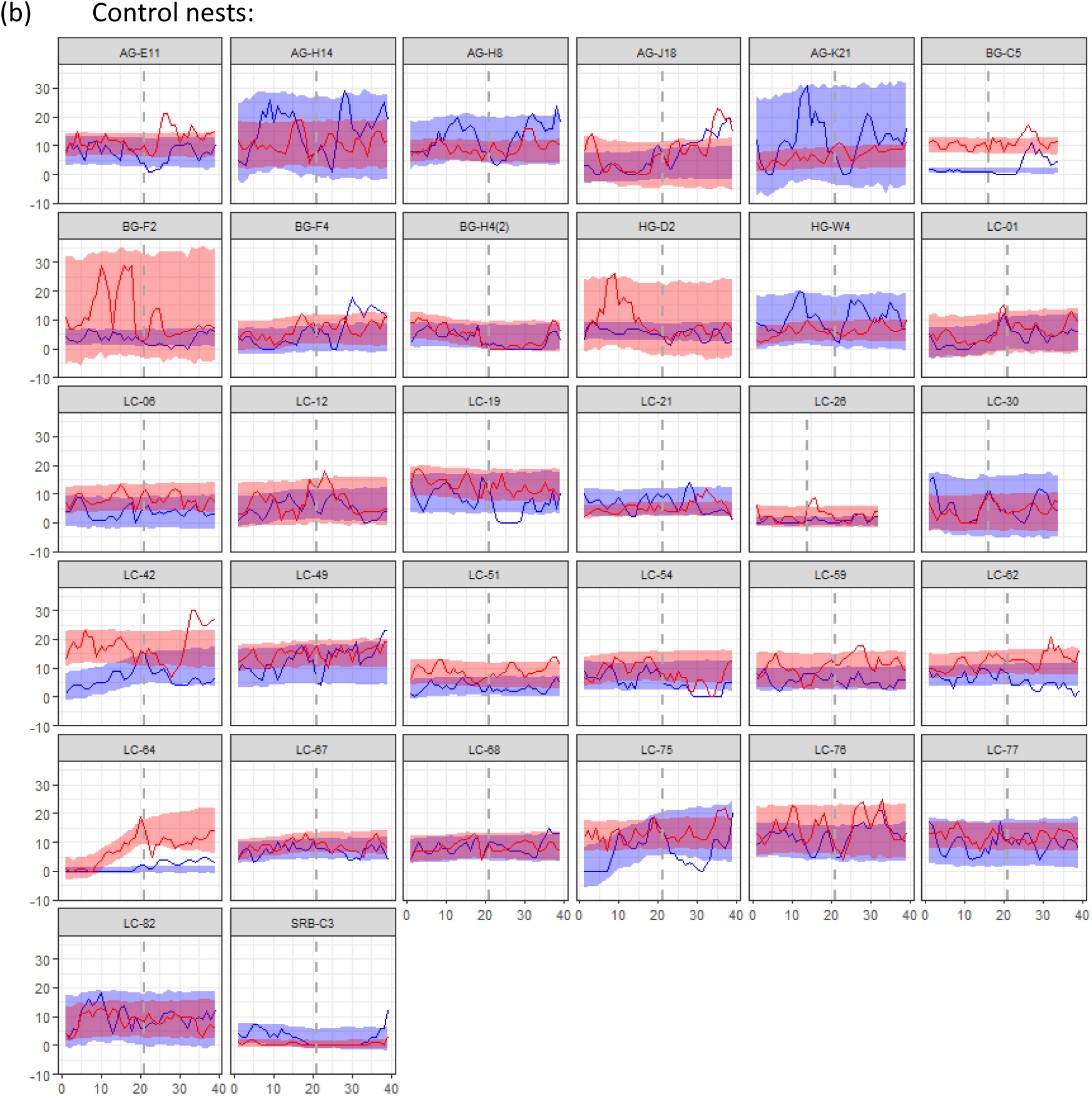
Behavioural coordination between male and female tree swallows. Blue and red colour represent males and females, respectively. The lines represent time series data of feeding visits, and the corresponding shaded areas denote 95% confidence range of the data observed during the pre-manipulation period (day 5) and prediction for the experimental period (day 6) using Bayesian structural time-series analysis (CausalImpact 1.2.1, Brodersen et al., Annals of Applied Statistics, 2015). The vertical dashed line indicates the start of the experimental period. When the lines go beyond the shaded area, it can be interpreted as a significant deviation from the prediction based on the pre-manipulation data.

**Figure S3:**
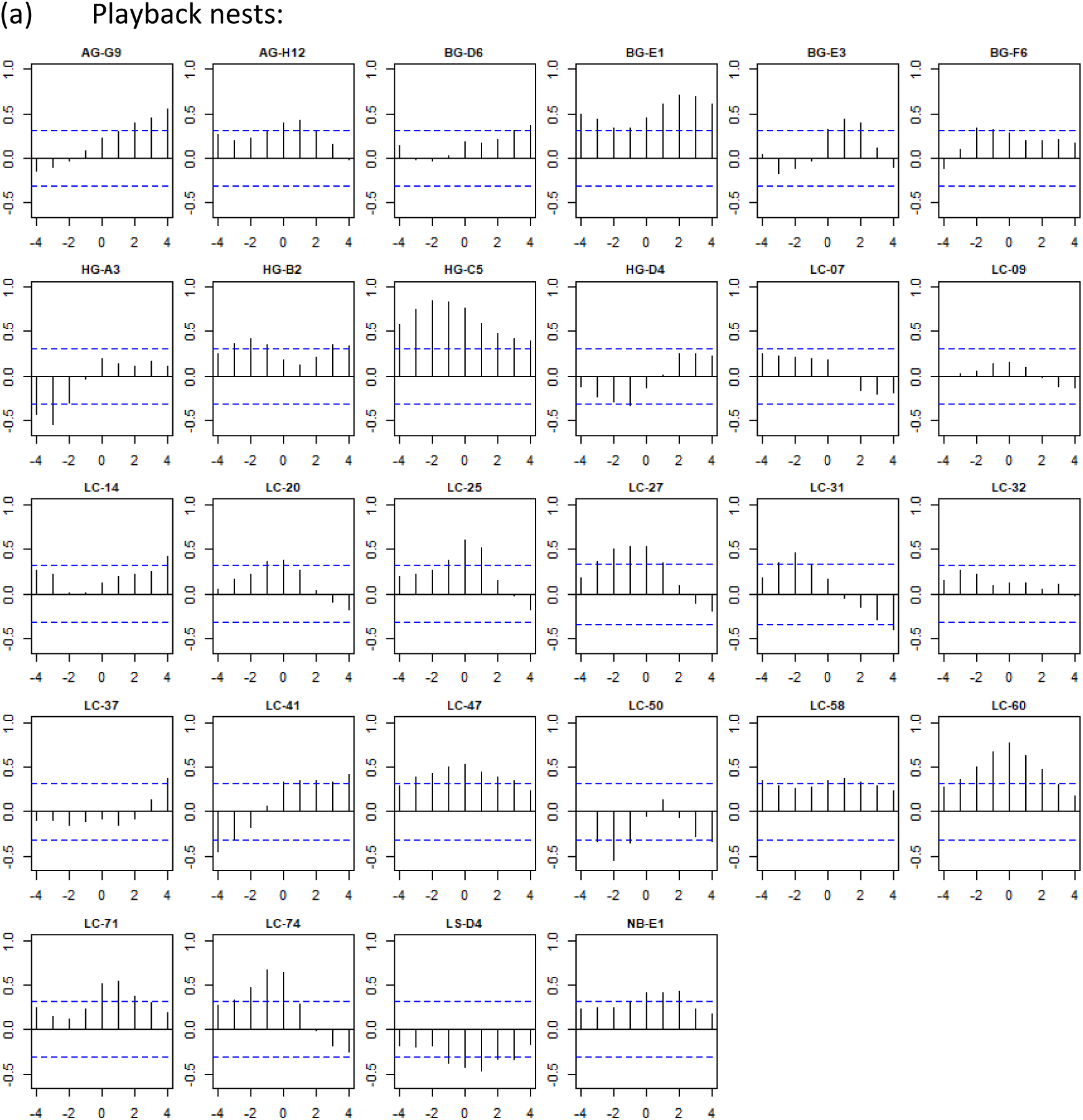
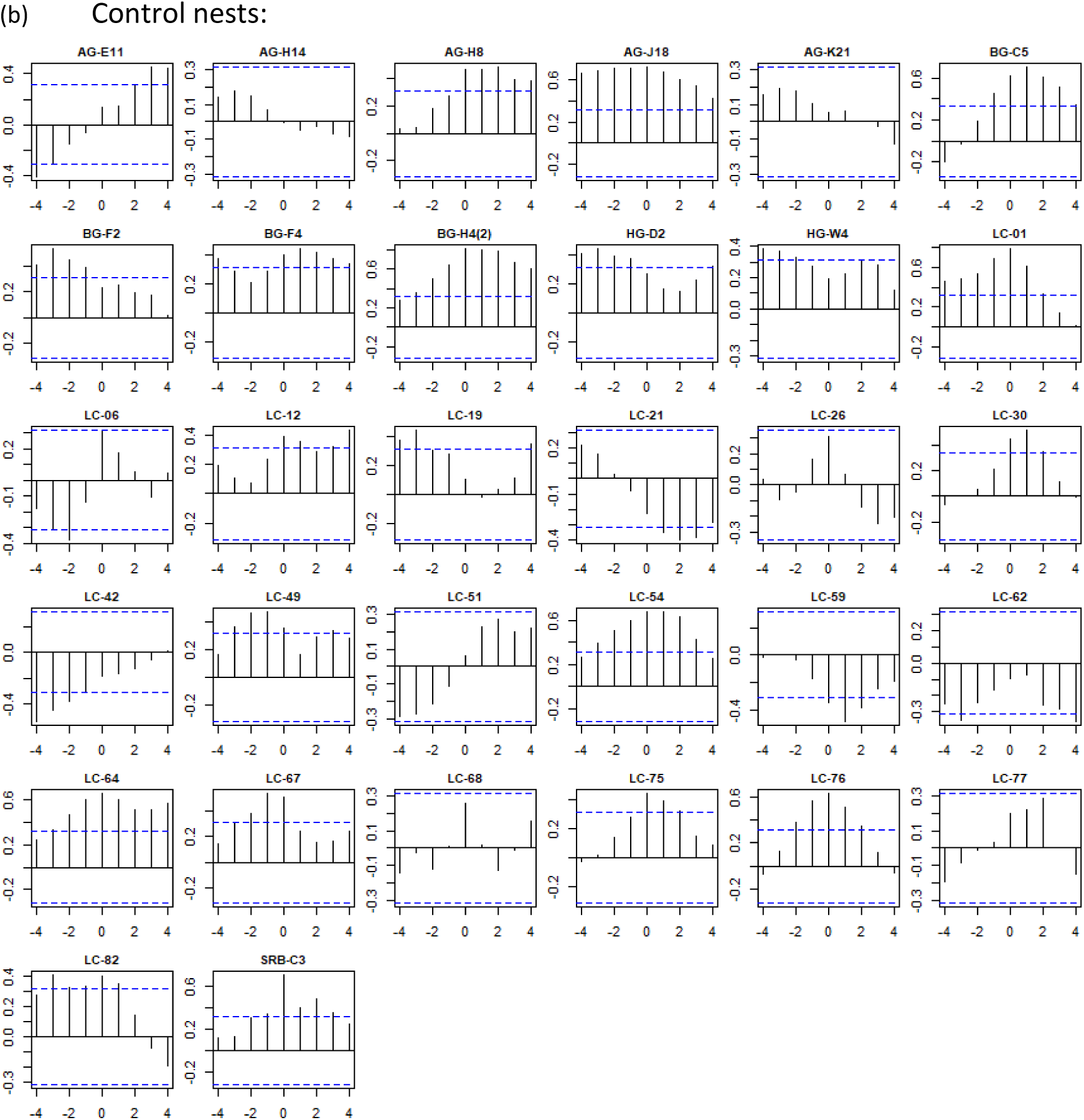
Cross-correlation coefficients of time series of female and male feeding visit behaviour in function of the lag (lag unit = 20 minutes). So when lag = 0, the pair is synchronous, where it is positive, then the covariation of the two time series is female-driven, where it is negative (i.e. males follow the females), where it is negative it is male-driven (females follow the males).

## The presence of males during playback broadcast to the females

Using 1h behavioural observations, we quantified the proportion of feeding visits of the females when their pair was present (in the vicinity of the nest box: either on the pole to which the nest box was attached or on the top of the nest box). The median proportion of female visits when the male was in the vicinity of the nest box was 0 in NC and 0.05 in Ontario. Males were significantly more often present during female visits in Ontario than in NC (p = 0.002), but the treatment did not affect their presence (p = 0.54) or the difference between the populations (p = 0.93). However, based on the RFID logs, males’ feeding visits in Ontario overlapped more often with the feeding visits of females than in NC (p = 0.04).

## Analysis of nestling begging rates

We analysed 241 feeding visits from 3 control and 6 playback nests. The average begging rate increased from day5 to day 6 in the case of both male and female visits (Fig. S4, Table S1).

**Figure S4:**
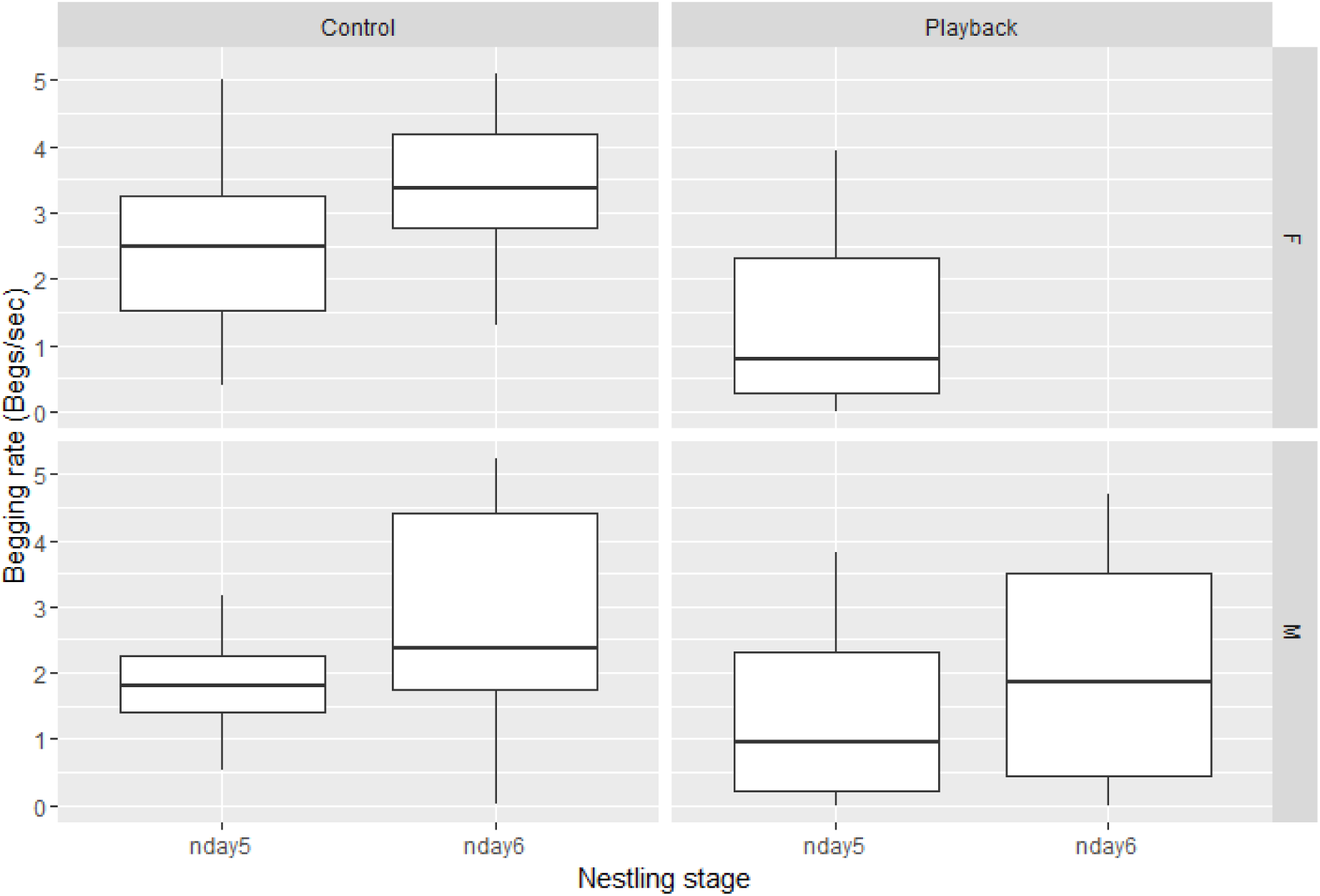
Nestling begging rate (number of begs per seconds) when male (M) or female (F) parents were feeding on day 5 (‘nday5’ – pre-treatment day) or day 6 (‘nday6’ – experimental playback period) post hatching.

**Table S1:** Parameter estimates for the model analysing nestling begging rate during male and female feeding visits on the pre-treatment day and during the experimental playback period (day 6).

| term | estimate | std. error | statistic | p-value |
| --- | --- | --- | --- | --- |
| **(Intercept)** | **1.74** | **0.34** | **5.18** | **<0.001** |
| **stage (day 6)** | **1.12** | **0.23** | **4.77** | **<0.001** |
| sex (male) | -0.15 | 0.17 | -0.87 | 0.385 |
| stage*sex (day6:male) | -0.26 | 0.31 | -0.83 | 0.409 |

Because of the overlapping playback sounds, the begging calls of the nestlings on day 6 for female visits could not be reliably detected, we could only analyse the interaction of the treatment and the nestling stage (day 5 vs. day 6) for male visits. In male visits alone, the increase in begging rate from day 5 to day 6 was still significant (p = 0.013), but the treatment did not affect the latter relationship (p = 0.732). In other words, nestlings in experimental nests did not increase their begging rate more than control nestlings, so the treatment did not affect the begging rate - at least towards males. This finding makes it unlikely that the significant increase in Ontario male feeding rates in response to the treatment is due to an increased begging rate of the experimental nestlings.

